# Transthyretin promotes axon growth via regulation of microtubule dynamics and tubulin acetylation

**DOI:** 10.1101/2021.03.24.436762

**Authors:** Jessica Eira, Joana Magalhães, Nídia Macedo, Maria Elena Pero, Thomas Misgeld, Mónica M Sousa, Francesca Bartolini, Márcia A Liz

**Affiliations:** ICBAS, Instituto de Ciências Biomédicas Abel Salazar, Universidade do Porto, Porto, Portugal; Neurodegeneration Team Instituto de Biologia Molecular e Celular- IBMC, and i3S – Instituto de Investigação e Inovação em Saúde, Universidade do Porto, Porto, Portugal; Nerve Regeneration Group, Instituto de Biologia Molecular e Celular- IBMC, and i3S – Instituto de Investigação e Inovação em Saúde, Universidade do Porto, Porto, Portugal; Department of Pathology & Cell Biology, Columbia University, New York; Department of Veterinary Medicine and Animal Production, University of Naples Federico II, Naples, Italy; Institute of Neuronal Cell Biology, Technical University of Munich, Munich, Germany

**Keywords:** Transthyretin, nerve biology, axon growth, microtubules, tubulin acetylation

## Abstract

Transthyretin (TTR), a plasma and cerebrospinal fluid protein, increases axon growth and organelle transport in sensory neurons. These TTR functions were suggested to underlie its activity in promoting nerve regeneration. While neurons extend their axons, the microtubule (MT) cytoskeleton is crucial for the segregation of functional compartments and axonal outgrowth. Herein, we investigated the hypothesis that TTR promotes axon elongation and regeneration by modulating MT dynamics. Indeed, we found that TTR KO mice have an intrinsic increase in dynamic MTs and reduced levels of acetylated α-tubulin in uninjured peripheral axons, and fail to modulate microtubule dynamics in response to sciatic nerve injury. Importantly, restoring acetylated α-tubulin levels of TTR KO DRG neurons using an HDAC6 inhibitor was sufficient to completely revert defective MT dynamics and neurite outgrowth. In summary, our results revealed a new role for TTR in the modulation of MT dynamics by regulating α-tubulin acetylation and support that this activity underlies TTR neuritogenic function.

## Introduction

Transthyretin (TTR) is a homotetrameric protein predominantly synthesized in the liver and the choroid plexus of the brain that functions as a thyroxin (Woeber & Ingbar, 1968) and retinol transporter (Goodman, 1984) in the plasma and cerebrospinal fluid. When mutated TTR aggregates and deposits in the peripheral nervous system (PNS), resulting in a familial form of peripheral neuropathy – familial amyloid neuropathy (Plante-Bordeneuve & Said, 2011). Besides causing neurodegeneration, TTR participates in nerve physiology and repair as inferred by the phenotype of TTR KO mice which display sensorimotor impairment and decreased regenerative capacity following sciatic nerve crush (Fleming, Saraiva, & Sousa, 2007). Further studies demonstrated that TTR is internalized by sensory dorsal root ganglia neurons (DRG), increasing neurite outgrowth and promoting axonal transport (Fleming, Mar, Franquinho, Saraiva, & Sousa, 2009). However, the cellular and molecular details underlying the neuritogenic activity of TTR remain to be deciphered.

Microtubules (MTs) play a fundamental role in neuronal health not only by providing structural support and establishing the tracks for axonal transport, but also as the modulation of MT dynamics is essential for proper axon growth and synaptic function, either during development or upon nerve regeneration. After injury, peripheral axons form a growth cone invaded by dynamic MTs while axon shafts contain more stable MT bundles (Erturk, Hellal, Enes, & Bradke, 2007). In addition to tubulin nucleotide binding state, tubulin isoforms and the action of MT-associated proteins (MAPs), tubulin post-translational modifications (PTMs) are important regulators of MT dynamics (Moutin, Bosc, Peris, & Andrieux, 2020). Among several tubulin PTMs, α-tubulin acetylation, a tubulin PTM associated with stable MTs, plays an essential role in mechanosensory neurons (Yan et al., 2018). Notably, increasing this PTM by inhibiting the enzyme responsible for deacetylation, HDAC6, promotes axon regeneration by increasing MT stability in the axon shaft (Rivieccio et al., 2009).

In this study, we investigated whether TTR neuritogenic activity in DRG neurons is conveyed by regulation of microtubule dynamics. Indeed, we found that in the course of axon growth, TTR increases MT dynamics in the distal end of growing axons while it stabilizes MT dynamics in the axon shaft. We further show that TTR promotes neurite outgrowth via regulation of tubulin acetylation in the axon shaft. Accordingly, the use of an HDAC6 inhibitor to rescue acetylated tubulin levels restores MT dynamics and neurite outgrowth of TTR KO neurons.

## Results and Discussion

### TTR induces compartmentalized modulation of microtubule dynamics in DRG neurons, to produce increased neurite outgrowth

TTR increases neurite outgrowth of DRG neurons through an unknown mechanism (Fleming et al., 2007). Herein we set out to determine whether TTR neuritogenic activity is promoted by its possible interference with MT dynamics. Soluble wild type (WT) TTR added to primary cultures of 4-week-old mouse DRG neurons promoted neurite outgrowth as determined by a significant increase in the length of the longest neurite (Figure 1 – figure supplement 1A-C). We tested whether this effect occurred with a concurrent modulation of MT dynamics and found that in the growth cone of DRG neurons transfected with the MT plus-tip binding protein EB3, WT TTR significantly increased EB3 growth rate and comet growth length, while it decreased duration of growth (Figure 1A-D, Video 1,2). Additionally, TTR increased the density of dynamic EB3 comets (Figure 1A, E). These results suggest that TTR promotes neurite outgrowth by increasing MT plus end dynamics in the growth cone of DRG neurons.

**Figure 1.**
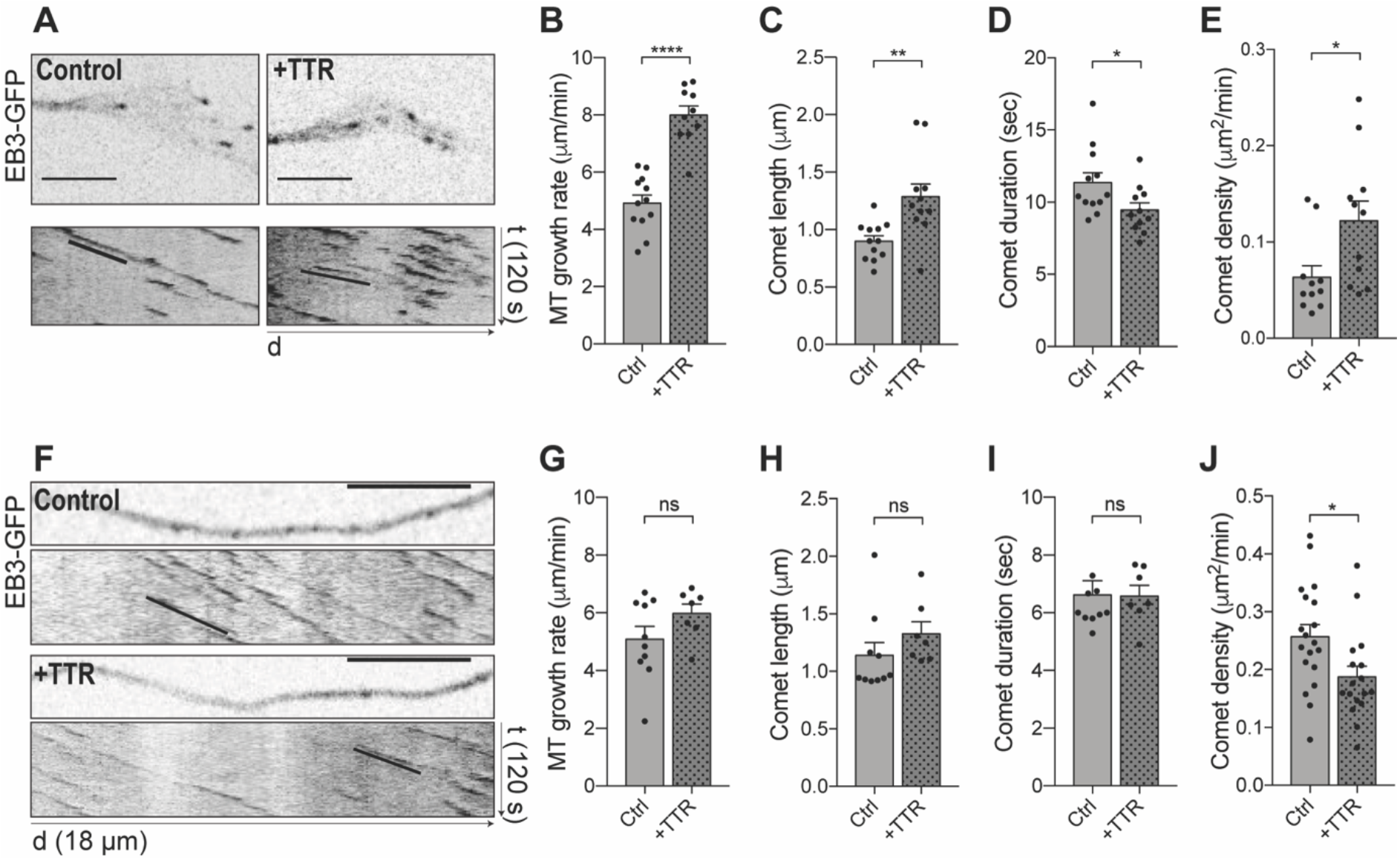
Soluble TTR has a compartmentalized impact on MT dynamics in DRG neurons. **(A)** Representative still images of the growth cones from DRG neurons transfected with EB3-GFP either untreated (Control) or treated with soluble WT TTR (+TTR) (top) and correspondent kymographs (bottom). **(B-E)** Quantifications of different MT dynamics parameters in the growth cone including MT growth rate (B), comet length (C), comet duration (D) and EB3 comet density (E), related to A. Results are plotted as mean ± SEM (n = 10-12 growth cones/condition, representative of 3 independent experiments). Statistical significance was determined by Student’s t-test: *P<0.05, **P<0.01, ****P<0.0001. Scale bars: 5 μm. **(F)** Representative still images of the neurite shafts from DRG neurons transfected with EB3-GFP either untreated (Control) or treated with WT soluble TTR (+TTR) and corresponding kymographs. **(G-J)** Quantifications of different MT dynamics parameters in the axon shaft including MT growth rate (G), comet length (H), comet duration (I) and EB3 comet density (J) related to F. Results are plotted as mean ± SEM (n = 7-19 neurite shafts/condition, representative of 3 independent experiments). Statistical significance was determined by Student’s t-test: *P<0.05; ns, not significant. Scale bars: 5 μm.

MT dynamicity is highly dependent on subcellular localization, especially in neurons, where MTs are not attached to the centrosome and MT plus ends are subjected to different stimuli depending upon whether their location is proximal or distal relative to the cell body. We investigated the effect of WT TTR on the dynamic state of MTs in the neurite shaft of DRG growing neurons and found that it had no impact on comet growth rate, comet length or in growth duration (Figure 1F-I). However, and in contrast to what was observed in the growth cone, TTR decreased EB3 comet density in the neurite shaft (Figure 1J). Of note, dynamic MT ends serve as a general marker of MT dynamics and of ongoing remodeling of axonal arbors (Kleele et al., 2014). In addition, while in the growth cone MTs must be highly dynamic to support growth and respond to extracellular stimuli during neurite extension, stable MTs are needed in the axon shaft to drive growth forward (Bradke, Fawcett, & Spira, 2012). Although further studies should explore how TTR promotes this compartmentalized effect, our data offer a mechanistic explanation of the pro-growth activity of TTR in axon extension by inducing differential regulation of MT dynamics.

### TTR KO sciatic nerve axons have increased MT plus ends and fail to increase MT dynamics in response to sciatic nerve crush *in vivo*

Based on the observation that TTR modulates MT dynamics *in vitro* while inducing axon outgrowth, we investigated whether genetic ablation of TTR may affect MT dynamics *in vivo* during nerve regeneration after sciatic nerve crush (Fleming et al., 2007). For that, we analysed *in vivo* MT dynamics using TTR KO-Thy1-EB3-YFP and their control WT-Thy1-EB3-YFP littermates (crossings are detailed in the methods section). We performed sciatic nerve crush in 12-week-old mice, and at the 3rd day post injury we collected both the ipsilateral (crushed) and the contralateral (uninjured) nerves prior to *ex vivo* live imaging of EB3 comets (Figure 2A). After crush, WT sciatic nerves mounted a response to injury characterized by increased comet growth rate and density distal to the crush site, when compared with uninjured nerves (Figure 2B, D, E, Video 3, 4). In contrast, injury to TTR KO mice, did not elicit an increase in comet growth speed or density (Figure 2C, D, E, Video 5, 6). While analysing uninjured sciatic nerves, we also observed that under basal conditions, TTR KO mice had higher EB3 comet density when compared to WT animals (Figure 2E). These results indicated that loss of TTR promoted either MT plus end rescue events or *de novo* MT nucleation. It was further conceivable that loss of TTR might upregulate levels of MT severing enzymes, a condition that would increase the number of growing MT plus ends by generating internal breaks in the MT polymer (Zhang, Fishel, Bertroche, & Dixit, 2013). To rule out these possibilities, we assessed the density of individual MTs by electron microscopy analysis of cross sections of WT and TTR KO sciatic nerves and measured the levels of MT severing enzymes on protein extracts from TTR KO and WT naïve sciatic nerves. Only axons with 2-4 μm were included in the electron microscopy analysis since those were also the ones that contributed to the differences in comet density observed by EB3 live-imaging. No difference was scored in MT density between WT and TTR KO axons (Figure 2F, G) and western blot analysis showed no fluctuation in the levels of both spastin and katanin on protein extracts from TTR KO and WT naïve sciatic nerves (Figure 2 – figure supplement 1A, B). These results indicated that in the absence of TTR, naïve axons in peripheral nerves do not nucleate more MTs or sever pre-existing ones but rather display an intrinsic increase in the frequency at which MT plus ends switch from depolymerization to polymerization (rescue), a condition that might account for their inability to reorganize the MT cytoskeleton after injury to regenerate. These results are in complete agreement with the increase in MT dynamics observed in the growth cones of growing neurons exposed to TTR *in vitro* (Figure 1) and demonstrate that in the absence of TTR the MT cytoskeleton fails to respond to nerve injury *in vivo*. We analysed the total levels of α- and βIII-tubulins and found no difference between WT and KO nerves, a result compatible with similar MT density scored between the two genotypes by electron microscopy analysis (Figure 2 – figure supplement 1C, D).

**Figure 2.**
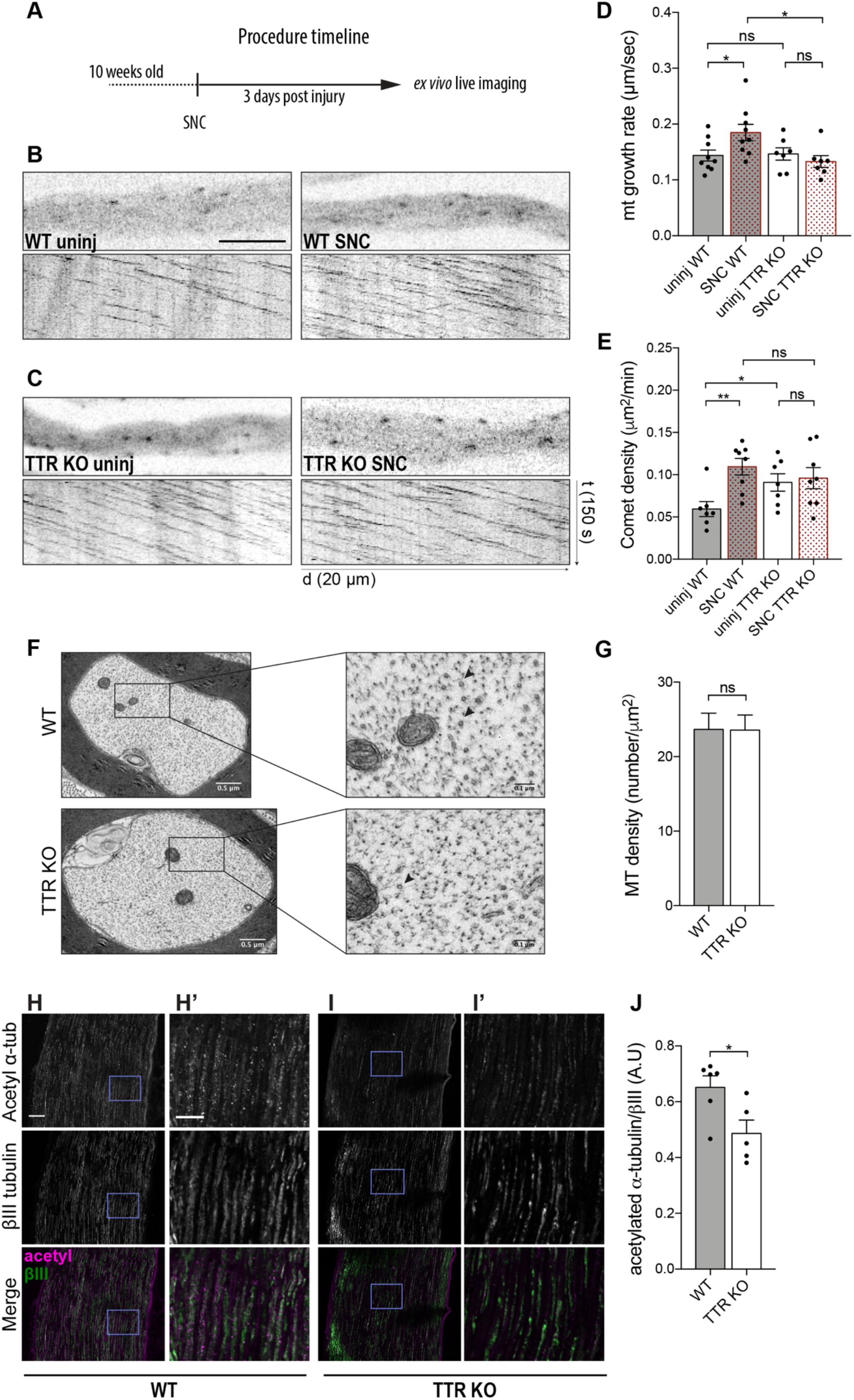
Sciatic nerve axons from TTR KO mice have more dynamic MT plus ends and fail to modulate MT dynamics in response to sciatic nerve crush *in vivo*. **(A)** Schematic representations of the procedure timeline of the experimental setup for MT dynamics assessment. **(B)** Representative still images of the axon shaft from a WT-Thy1-EB3-YFP uninjured nerve (WT uninj) with correspondent kymograph and of an axonal region distal to the lesion site from a WT-Thy1-EB3-YFP crushed nerve (WT SNC) with correspondent kymograph. Scale bar: 5 μm. **(C)** Representative still images of the axon shaft from a TTR KO-Thy1-EB3-YFP uninjured nerve (TTR KO uninj) with correspondent kymograph and of an axonal region distal to the lesion site from a TTR KO-Thy1-EB3-YFP crushed nerve (TTR KO SNC) with correspondent kymograph. **(D, E)** Quantifications of different MT dynamics parameters including MT growth rate (D) and EB3 comet density (E) from WT-Thy1-EB3-YFP (WT) and TTR KO-Thy1-EB3-YFP (TTR KO) nerves either uninjured (uninj) or crushed (SNC). Results are plotted as mean ± SEM (n = 7-9 animals/condition, 6-12 axons/animal). Statistical significance determined by Student’s t-test: *P<0.05, **P<0.01. ns, not significant. **(F)** Representative electron microscopy images of individual axons from 10-week-old WT and TTRKO sciatic nerves and respective high magnification representations. MTs are represented by black arrowheads. **(G)** Quantification of MT density from 2-4 um axons from WT and TTR KO sciatic nerves. Results are plotted as mean ± SEM (n = 6 animals/genotype, 30-60 axons/animal). Statistical significance determined by Student’s t-test: ns, not significant. **(H, I)** Representative images of 10-week-old WT (H) and TTR KO (I) sciatic nerves immunostained for acetylated α-tubulin and βIII-tubulin. Scale bar: 50 μm. (H’, I’) Zoomed in regions from H and I, respectively. Scale bar: 20 μm. **(J)** Quantification of the relative values of acetylated α-tubulin over βIII-tubulin. Data represent mean ± SEM (n = 5-6 animals/genotype, 45-57 axons/animal). Statistical significance was determined by Student’s t-test: *P<0.05.

Acetylated α-tubulin accumulates onto previously stabilized MTs but its presence has been further associated to an increase in MT longevity given the enhanced ability of acetylated MTs to resist breakage (Xu et al., 2017). In addition, recent studies have shown that loss of acetylated α-tubulin leads to increased axonal MT debundling and MT plus-end dynamics with consequences in CNS development (Dan et al., 2018). Given the impact of acetylated α-tubulin levels on axonal integrity and sensory neuron function, we analysed this PTM by immunohistochemistry and found that the ratio of acetylated α-tubulin versus βIII-tubulin intensity on labelled axons was decreased in the sciatic nerves of TTR KO mice (Figure 2H-J). Altogether, our data strongly support the notion that loss of acetylated α-tubulin in TTR KO mice may lead to an increase in MT dynamicity and suggest that in TTR KO mice these defects contribute to the inability of their nerves to regenerate.

TTR KO mice also display a sensorimotor impairment that starts at 6 months of age (Fleming et al., 2007). In mouse peripheral sensory neurons, acetylated α-tubulin is enriched in submembranous bands in the soma rather than in the cytoplasmic MT network (Morley et al., 2016) and this somatic enrichment is thought to tune the mechanical properties of the membrane. Indeed, neurons deprived of acetylated tubulin are less elastic and require more force to trigger the mechanosensitive channels, a property corroborated by the importance of acetylated α-tubulin in maintaining touch sensitivity in mechanosensory neurons (Yan et al., 2018). This evidence along with our observations strongly suggest that sensorimotor defects in TTR KO mice might be related to dysregulation of α-tubulin acetylation. Additionally, TTR modulation of MTs might also underlie axonal transport impairment reported in TTR KO mice (Fleming et al., 2009).

### TTR promotes neurite outgrowth via regulation of tubulin acetylation in the axonal shaft

To investigate whether loss of acetylated α-tubulin in TTR KO nerves underlies the increase in the number of dynamic MT ends, we evaluated the consequences of exposing WT and TTR KO DRG neurons to the HDAC6 inhibitor ACY-738 to increase tubulin acetylation levels. To this end, DIV4 DRG neuron cultures isolated from adult WT and TTR KO mice were carried out and levels of acetylated tubulin were evaluated in these neurons by immunofluorescence. First, we confirmed that the neurite shafts of TTR KO had decreased levels of acetylated α-tubulin when compared to WT controls (Figure 3A, B, E). More importantly, while the addition of ACY-738 did not affect WT neurons, as it was used at a low concentration (Benoy et al., 2017), it normalized loss of α-tubulin acetylation in the neurite shafts from TTR KO neurons (Figure 3C, D, E). We analysed whether the increase in α-tubulin acetylation promoted by ACY-738 in TTR KO axons had an impact on the number of dynamic MTs by measuring EB3 comet density in WT-Thy1-EB3-YFP and TTR KO-Thy1-EB3-YFP neurons. Indeed, EB3 live imaging confirmed an increase in comet density in TTR KO neurite shafts when compared to WT controls (Figure 3F), similarly to what was observed *in vivo*. However, this effect was completely reverted by ACY-738 (Figure 3F), strongly supporting that regulation of MT dynamicity by TTR is mediated by its modulation of α-tubulin acetylation levels.

**Figure 3.**
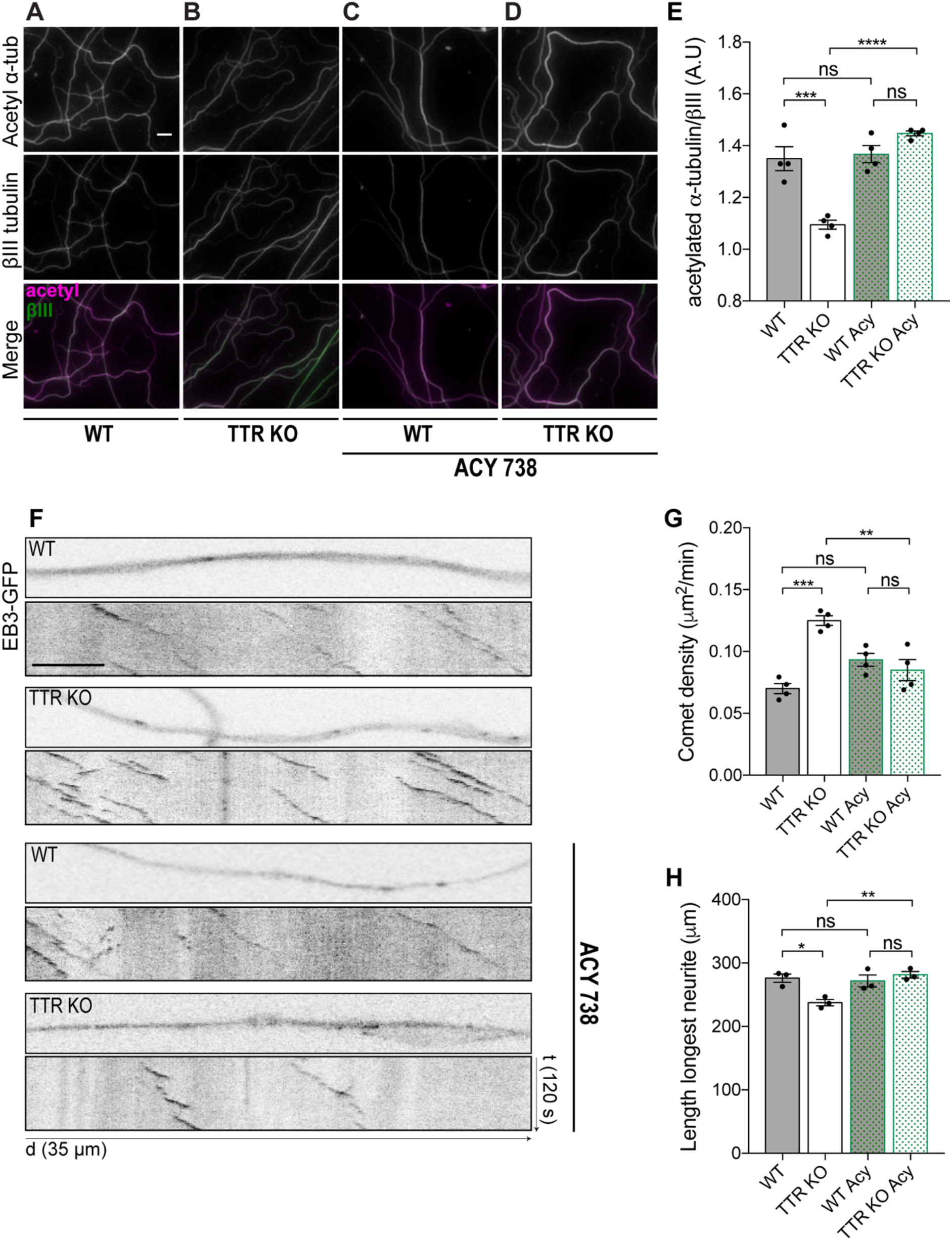
TTR-mediated regulation of MT dynamics and axonal growth occurs via modulation of tubulin acetylation. **(A-D)** Representative images of acetylated α-tubulin (magenta) and βIII-tubulin (green) immunostaining of WT and TTR KO 4DIV DRG cultures untreated (A, B) and treated with ACY-738 (C, D) Scale bar: 10 μm. **(E)** Quantification of the relative values of acetylated α-tubulin over βIII-tubulin. Data represent mean ± SEM (n = 4 independent samples/condition; 50 axons/sample). Statistical significance determined by Tukey’s multiple comparisons test: ***P<0.001, ****P<0.0001, ns - not significant. **(F)** Representative still images of the axonal shafts from 4DIV WT-Thy1-EB3-YFP (WT) and TTR KO-Thy1-EB3-YFP (TTR KO) DRG neurons either untreated or treated with ACY-738 and correspondent kymographs. Scale bar: 5 μm. **(G)** Quantification of comet density in the neurite shafts from DIV4 WT-Thy1-EB3-YFP and TTR KO-Thy1-EB3-YFP DRG neurons untreated (WT; TTR KO) or treated with ACY-738 (WT Acy; TTR KO Acy). Data represent mean ± SEM (n = 4, independent samples/condition. 9-17 neurite shafts/sample). Statistical significance determined by Tukey’s multiple comparisons test: **P<0.01 ***P<0.001, ns - not significant. **(H)** Quantification of the length of the longest neurite in DIV1 WT and TTR KO DRG neurons untreated (WT; TTR KO) or treated with ACY-738 (WT Acy; TTR KO Acy). Data represent mean ± SEM (n = 3 independent samples/condition. 50-114 neurons/sample). Statistical significance determined by Tukey’s multiple comparisons test: *P<0.05, **P<0.01, ns - not significant.

To ultimately demonstrate that TTR increases neurite outgrowth by interfering with MT dynamics, we assessed the effect of ACY-738 on the impaired neurite outgrowth of TTR KO neurons. Strikingly, while addition of ACY-738 did not affect neurite outgrowth of WT neurons, it successfully rescued the ability of TTR KO DRG neurons to extend neurites *in vitro* (Figure 3G). These data demonstrate that TTR regulation of tubulin acetylation levels is required for its neuritogenic activity.

Overall, our results reveal a novel activity of TTR as a regulator of MT dynamics by modulating acetylated α-tubulin levels, and demonstrate that this activity underlies the TTR-mediated promotion of axonal outgrowth. Moreover, we suggest that TTR modulation of α-tubulin acetylation might be related to the reduced axonal transport and sensorimotor impairment observed in TTR KO mice. A loss of TTR function may occur in familial amyloid polyneuropathy contributing to sensory neuropathy. Further studies are needed to decipher how TTR modulates acetylated α-tubulin levels in axons. TTR-dependent modulation of expression levels, localization and/or activities of the tubulin acetylase ATAT1 and the deacetylases HDAC6 and Sirtuin2, all involved in the tubulin acetylation cycle, are exciting areas of investigation in the future.

## Materials and Methods

### Animals

Mice were handled according to European Union and National rules. WT and TTR KO (Episkopou et al., 1993) littermates (in the Sv/129 background), were obtained from the offspring of heterozygous breeding pairs. Thy1.EB3-YFP (Kleele et al., 2014) were crossed with TTR KO mice. The resultant TTR KO(+/−).Thy1.EB3-YFP were intercrossed generating WT.Thy1.EB3-YFP and TTR KO.Thy1.EB3-YFP. All animals were maintained under a 12 h light/dark cycle and fed with regular rodent’s chow and tap water *ad libitum*. Genotypes were determined from ear extracted genomic DNA.

### Recombinant TTR production and purification

Recombinant WT TTR was produced in *Escherichia coli* BL21(DE3) cells transformed with a pETF1 vector carrying human WT TTR (Goldsteins et al., 1997). TTR protein was expressed as described in (Silva et al., 2017) and purified through successive ionic exchange, hydrophobic interaction and size exclusion chromatography. For cellular assays, recombinant TTR was detoxified using a high-capacity endotoxin removal resin (Thermo Scientific) and quantified using the Lowry based DC Protein Assay (Bio-Rad Laboratories).

### Primary DRG neuronal cultures and cell treatment

Primary cultures of DRG neurons from 4-to 8-week-old mice were performed as described in (Liz et al., 2014). For neurite outgrowth experiments with TTR addition, WT DRG neurons were plated in 20 μg/ml PLL + 5 μg/ml Laminin coated 24 well plates at a density of 5000 cells per well in complete medium (DMEM/F12 (Sigma-Aldrich, D8437) supplemented with 1× B27 (Gibco), 1% penicillin/streptomycin (Gibco), 2 mM L-glutamine (Gibco), and 50 ng/mL NGF (Millipore, 01-125)) at 37ºC and 5% CO_2_. 4 h after plating, DRG were treated with recombinant WT TTR (equal volume of PBS for the control condition) at a concentration of 300 μg/ml and incubated for 12 h at 37ºC and 5% CO_2_. For EB3-GFP transfection experiments, the 4D Nucleofector Amaxa system (Lonza, Barcelona, Spain, CM#137 program) was used and WT DRG neurons were nucleofected at a density of at least 200.000 cells/condition with a truncated version of EB3-GFP (a construct containing aminoacids 1 to 200 of EB3, artificially dimerized by the addition of the leucine zipper domain of GCN4, cloned into the pEGFP-N1 vector, that efficiently accumulates at microtubule tips) (Komarova et al., 2009). After transfection, cells were left in suspension for 24 h and then plated on 20 μg/ml PLL + 5 μg/ml Laminin coated 35 mm μ-dishes (iBidi) at a density of 15000 neurons per dish in phenol-free DMEM/F12 with its supplementation. 4 h after plating, DRG neurons were treated with recombinant WT TTR at a concentration of 300 μg/ml and incubated for 12 h at 37ºC and 5% CO_2_. For acetylated α-tubulin levels assessment and MT dynamics evaluation through HDAC6 inhibition using ACY-738 (provided by Acetylon Pharmaceuticals; (Jochems et al., 2014), WT.Thy1.EB3-YFP and TTR KO.Thy1.EB3-YFP DRG neurons were used. Neurons were plated on 20 μg/ml PLL + 5μg/ml Laminin coated 24 well plates at a density of 5000 cells per well for acetylated α-tubulin and βIII-tubulin staining or in 8 well μ-dishes (iBidi) at a density of 8000 cells per well for EB3 live imaging of MT dynamics, in DMEM/F12 (Sigma-Aldrich, D8437) supplemented with 1× B27 (Gibco), 1% penicillin/streptomycin (Gibco), 2 mM L-glutamine (Gibco), 50 ng/mL NGF (Millipore, 01-125), 60μM 5-Fluoro-2’-deoxyuridine (FluoU) and 100 nM of ACY-738 in DMSO (equal volume of DMSO for the control condition) at 37ºC and 5% CO_2_. At DIV3 half the medium was changed to new medium supplemented with 2× the concentration of ACY-738 and FluoU. At DIV 4, 2 h before fixing, medium was changed supplemented with ACY-738. For live imaging experiments, 2 h before imaging, medium was changed to phenol-free DMEM/F12 with its supplementation and imaging was started in the ACY-738 treated conditions to minimize the effect of its short life in plasma (Jochems et al., 2014). For neurite outgrowth assessment with HDAC6 inhibition using ACY-738, 10-week-old WT and TTR KO DRG neurons were plated on 20 μg/ml PLL + 5μg/ml Laminin coated 24 well plates at a density of 5000 cells per well in complete medium supplemented with 100 nM of ACY-738 in DMSO (equal volume of DMSO for the control condition) at 37ºC and 5% CO_2_ for 16 h.

### Sciatic nerve crush

12-week-old mice were anesthetized with isoflurane and a 4-mm-long incision was made in the shaved thigh skin. For nerve crush, the sciatic nerve was exposed and crush was performed using Pean forceps, three times during 10 s. To standardize the procedure, the crush site was maintained constant for each animal at 5 mm distally to the notch. A single skin suture, immediately above the crush site, served as an additional reference. After surgery, animals were allowed to recover for 3 days after which, mice were sacrificed in the CO_2_ chamber and the sciatic nerves (injured (SNC) and contralateral naïve (uninj) nerves) were subsequently collected for live cell imaging.

### EB3 live imaging for the analysis of MT dynamics

For analysis of MT dynamics, DRG neurons were recorded for 2 min (60 frames total) in phenol-free DMEM/F12 supplemented as mentioned above, at 37°C and 5% CO_2_, using a Spinning Disk Confocal System Andor Revolution XD with an iXonEM+ DU-897 camera and an IQ 1.10.1 software (ANDOR Technology). For the *ex vivo* imaging, sciatic nerves from WT.Thy1.EB3-YFP and TTR KO.Thy1.EB3-YFP were collected 3 days post injury and placed in 35 mm μ-Dish (iBidi) with phenol-free DMEM/F12, and recordings were performed as described above. For the quantification of the different EB3 dynamics parameters, kymographs were made using the Fiji KymoResliceWide plugin (distance, x axis; time, y axis). Starting and end positions of the traces were defined using the Fiji Cell Counter plugin generating comet length, which is the comet movement length in micrometres, comet duration, which is the comet lifetime in seconds, and growth rate, which is the comet length/comet duration. Comet density at the growth cone is the number of comets in 30 consecutive frames divided by the number of quantified frames and growth cone area. Comet density at the axonal shaft is the number of comets per micrometre squared per minute.

### Sciatic nerve ultrastructure preparation and MT density analysis

For ultrastructure analysis of MT density, 12-week-old WT and TTR KO mice were sacrificed using a CO_2_ chamber and sciatic nerves were collected and fixed in 4% glutaraldehyde in 0.1 M sodium cacodylate buffer (pH 7.4) for 5 days and processed for ultrathin sections as previously described (da Silva et al., 2014). For MT density analysis, 30-60 axons of 2-4 μm of diameter were analysed using Photoshop CS3 for image processing and mounting.

### Immunoblotting

Protein lysates from 12-week-old quick frozen sciatic nerves from WT and TTR KO mice were prepared in ice-cold RIPA lysis buffer (1% Triton X-100, 0.1% SDS, 140 mM NaCl, 1× TE pH 8, 1× protease inhibitor Cocktail and 1 mM Sodium orthovanadate), sonicated (2×10 cycles, Output Power 50 Watts, Branson sonifier 250) and cleared by centrifugation at 15000 rpm for 10 min at 4^°^C. 20 ug of protein extracts for analysis of MT severing enzymes or 2.5 ug of protein extracts for analysis of tubulin proteins were separated under denaturing conditions and transferred to Amersham Protran Premium 0.45 μm nitrocellulose membranes (GE Healthcare Life Sciences) prior to blocking in 5% non-fat dried milk in TBS-T for 1 h at room temperature. Membranes were probed overnight at 4^°^C with the following primary antibodies (in 5% BSA in TBS-T): rabbit anti-katanin (1:500, Proteintech, 17560-1-AP), mouse anti-spastin (1:500, Santa Cruz Biotechnology, sc-81624), mouse anti-βIII-tubulin (1:10,000; Promega, G7121), mouse anti-α-tubulin (0.5 μg/ml; DSHB, 12G10), mouse anti-GAPDH (1:1,000; Santa Cruz Biotechnology, sc-166574) and rabbit anti-vinculin (3:10,000; ThermoFisher Scientific, 700062). Secondary antibodies were used in 5% non-fat dried milk in TBS-T for 1 hour at room temperature. Secondary antibodies were mouse IgGκ light chain conjugated with horseradish peroxidase (HRP) (1:2,000; Santa Cruz Biotechnology, sc-516102) and goat anti–rabbit IgG conjugated with HRP (1:10,000; Jackson ImmunoResearch Labs, 111-035-003). Immunodetection was performed by chemiluminescence using ECL (Millipore, WBLUR0500) and quantified using ImageJ software.

### Immunohistochemistry

Sciatic nerves from 12-week-old from WT and TTR KO mice were perfused with PBS for 5 min followed by 4% paraformaldehyde (PFA, pH 7.4) in PBS (40 ml). Sciatic nerves were collected and maintained in 4% PFA for 24 h and then cryopreserved in 30% sucrose for 48 h. Cryopreserved sciatic nerves were embedded in Optimum Cutting Temperature (OCT) compound (ThermoFisher Scientific), frozen and cut longitudinally (Cryostat Leica CM3050S) in 12 μm thick sections. For acetylated α-tubulin and βIII-tubulin staining, sections were blocked with 5% normal donkey serum containing 0,3% Triton in PBS for 1 h at RT followed by incubation for two overnights at 4^°^C with mouse anti-acetylated α-tubulin (1:500, Sigma-Aldrich, T7451) and rabbit anti-βIII-tubulin (1:500, Abcam, 1967-1) diluted in blocking buffer. Sections were then incubated with donkey anti mouse Alexa Fluor 568 (1:1000, Alfagene, A10037) and donkey anti rabbit Alexa Fluor 647 (1:500, Jackson ImmunoResearch, 711-605-152) diluted in blocking buffer for 1 h at RT, washed in PBS and mounted in iBidi mounting medium (iBidi, 50001).

### Immunocytochemistry

For neurite outgrowth experiments, neurons were fixed with 4% PFA, blocked with 5% normal donkey serum (NDS) containing 0,4%Tween for 1 h at RT, incubated with mouse anti–βIII-tubulin (1:2000; Promega, G7121) overnight at 4ºC followed by incubation with secondary antibody (donkey anti-mouse Alexa-Fluor 488, 1:1000 Alfagene, A21202) for 1 hr at RT. For acetylated α-tubulin and βIII-tubulin staining, WT DRG and TTR KO either untreated or treated with ACY-738, were fixed with cytoskeleton preservation PHEM fixative (4% PFA, 4% sucrose, 0.25% Glutaraldehyde, 0.1% Triton X-100, 300 mM PIPES, 125 mM HEPES, 50 mM EGTA and 10 mM Magnesium Chloride), permeabilized with 0.2% Triton X-100 for 5 min, quenched with 50 mM NH_4_Cl for 5 min and blocked with 2% Fetal Bovine Serum (FBS), 2% BSA and 0.2% Fish Gelatin in PBS for 1 h at RT. Incubation with mouse anti acetylated α-tubulin (1:5000; Sigma-Aldrich, T7451) and rabbit anti-βIII-tubulin (1:500, Abcam, 1967-1) was performed in 10% blocking buffer overnight at 4ºC. Incubation of the secondary antibodies donkey anti mouse Alexa Fluor 568 (1:1000, Alfagene, A10037) and donkey anti rabbit Alexa Fluor 647 (1:500, Jackson ImmunoResearch, 711-605-152) was performed in 10% blocking buffer for 1 h at RT. Cells were mounted in Fluoromount (Southern Biotech).

### Imaging and quantification

For neurite outgrowth experiments, images were acquired either using an epifluorescence microscope Zeiss AxioImager Z1 with an Axiocam MR3.0 camera and Axiovision 4.7 software or a Leica DMI6000 FFW microscope with an HC PL Fluotar 10× objective and a LAS X software operating the navigator feature. The longest neurite tracing analysis was performed using Fiji software and neuronJ plugin. For acetylated α-tubulin/βIII-tubulin quantifications in sciatic nerves, WT and TTR KO sections were imaged in a laser scanning Confocal Microscope Leica TCS SP8, using the PL APO 10x objective. For acetylated α-tubulin/βIII-tubulin quantifications in DRG neurons, Leica DMI6000 FFW microscope with an HCX PL Fluotar 20× objective was used with LAS X software. Ratiometric analysis of acetylated α-tubulin/βIII-tubulin in sciatic nerve slices and DRG neurons was performed by determination of regions of interest selecting stretches of axons, randomly, and using ImageJ software.

### Statistics

All statistical tests were performed using GraphPad Prism 7 software. Data are shown as mean ± s.e.m. Unless otherwise stated, the Student’s t test was used. Statistical tests and sample sizes are indicated in figure legends and significance was defined as *p < 0.05; **p < 0.01; ***p < 0.001; ****p < 0.0001; ns - not significant.

## Acknowledgements

This work was supported by: FEDER - Fundo Europeu de Desenvolvimento Regional funds through the COMPETE 2020 – Operacional Programme for Competitiveness and Internationalisation (POCI), Portugal 2020, and by Portuguese funds through FCT - Fundação para a Ciência e a Tecnologia/Ministério da Ciência, Tecnologia e Ensino Superior in the framework of the project POCI-01-0145-FEDER-028336 (PTDC/MED-NEU/28336/2017); Norte-01-0145-FEDER-000008 - Porto Neurosciences and Neurologic Disease Research Initiative at I3S, supported by Norte Portugal Regional Operational Programme (NORTE 2020), under the PORTUGAL 2020 Partnership Agreement, through FEDER; and Thompson Family Foundation (TFFI) award, RO1AG050658 (NIH/National Institute on Aging) and R21NS120076 (NIH/NINDS) awards to FB, and a PRIN-2017FJC3-004 (MIUR) to MEP. JE is a FCT fellow (SFRH/BD/116343/2016). MAL is an FCT Investigator (IF/00902/2015).

## Author contributions

MAL coordinated research. JE and MAL conceived and analysed the experiments. JE, JM, NM and MEP performed experiments. TM and MMS provided critical tools and revised the manuscript. JE, FB and MAL wrote the manuscript. All authors read and approved the final manuscript.

## Ethics

Animal Experimentation: All animals were handled according to the European Union Directive 2010/63/EU as well as the national Decreto-lei nº113-2013. The protocols described in this work have been approved by the IBMC/i3S 593 Ethical Committee and by the Portuguese Veterinarian Board.

## Supplementary information

**Figure S1.**
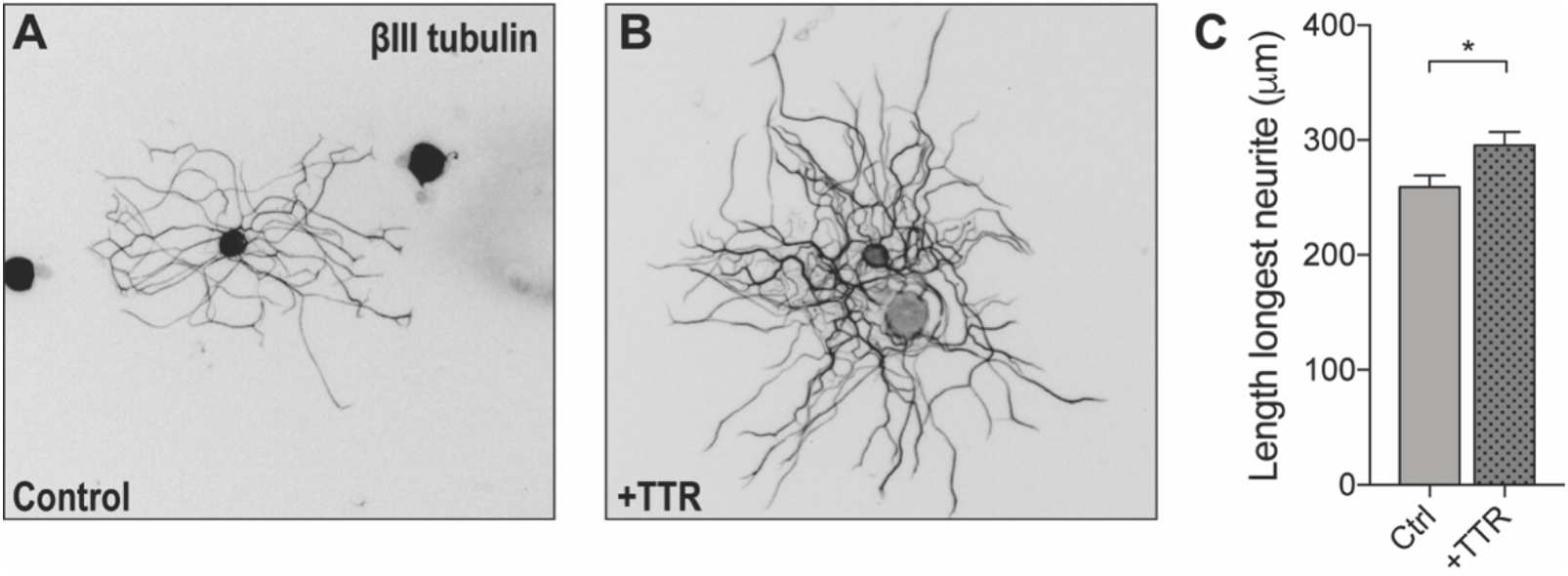
Exogenous TTR addition to DRG neurons increases neurite outgrowth. Representative anti-βIII-tubulin immunofluorescence images of control **(A)** and WT TTR treated **(B)** DRG neurons. **(C)** Quantification of the length of the longest neurite in DRG neurons treated with TTR. Data represent mean ± SEM (n = 112-197 neurons/condition, representative of 3 independent experiments). Statistical significance determined by Student’s t-test: *P<0.05.

**Figure S2.**
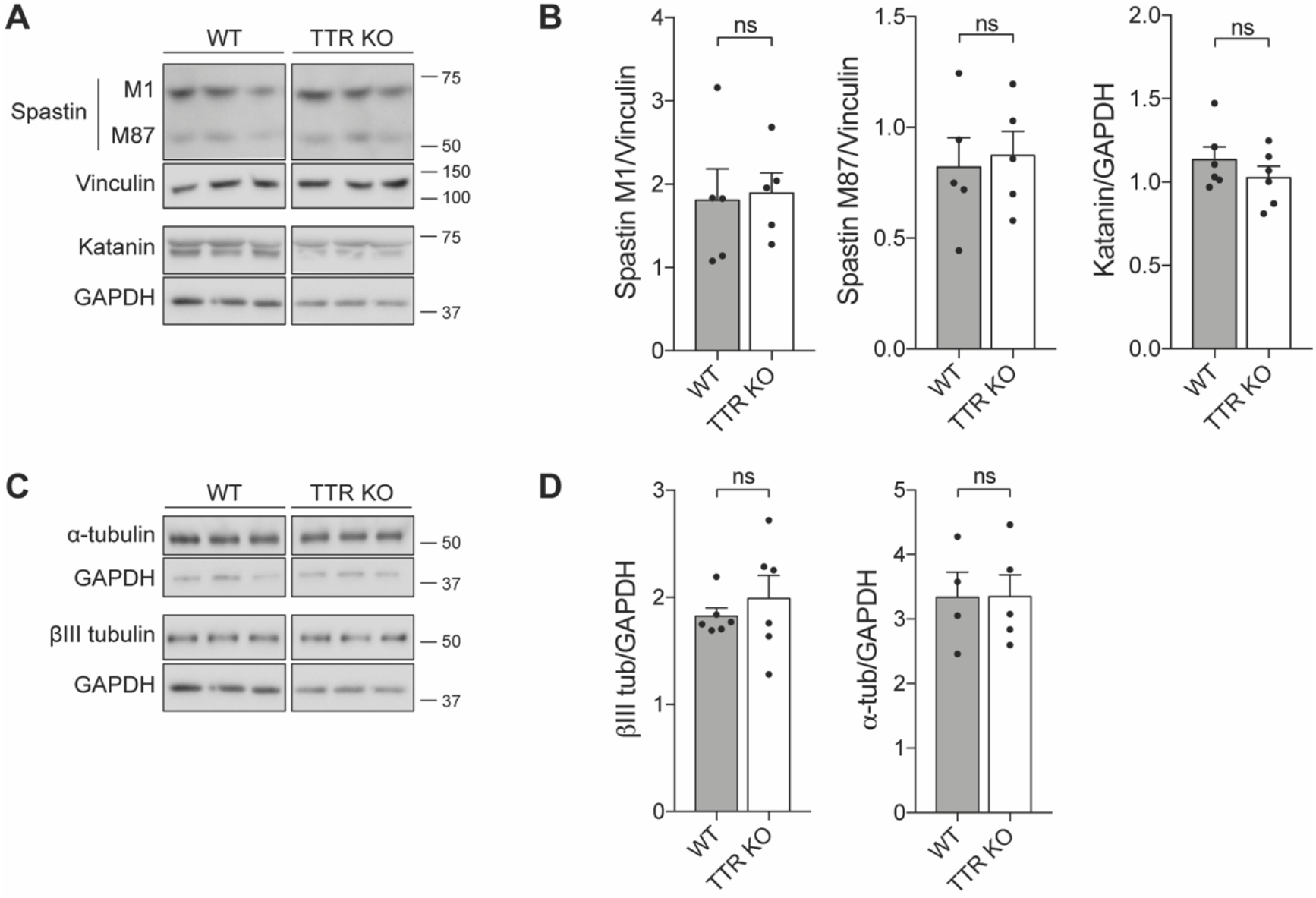
TTR KO animals display normal values of MT severing enzymes and total α- and βIII-tubulin. **(A, B)** Representative western blot (A) and respective quantification (B) showing MT severing enzymes levels in sciatic nerves of 12-week-old WT and TTR KO mice. Vinculin was used as control for Spastin and GAPDH was used as a control for Katanin. Data represent mean ± SEM (n = 5-6 animals/condition). ns – not significant by Student’s t test. **(C, D)** Representative western blot (C) and respective quantification (D) showing α- and βIII-tubulin levels in sciatic nerves of 12-week-old WT and TTR KO mice. GAPDH was used as a control. Data represent mean ± SEM (n = 4-6 animals/condition). ns - not significant by Student’s t test.

---

**Video 1. Growth cone from a control DRG neuron transfected with EB3-GFP.**

**Video 2. Growth cone from a DRG neuron transfected with EB3-GFP treated with WT soluble TTR.**

**Video 3. Axonal shaft from a WT-Thy1-EB3-YFP uninjured nerve.**

**Video 4. Axonal region distally to the lesion site from a WT-Thy1-EB3-YFP crushed nerve.**

**Video 5. Axonal shaft from a TTR KO-Thy1-EB3-YFP uninjured nerve.**

**Video 6. Axonal region distally to the lesion site from a TTR KO-Thy1-EB3-YFP crushed nerve.**

## References

Benoy, V., Vanden Berghe, P., Jarpe, M., Van Damme, P., Robberecht, W., & Van Den Bosch, L., (2017). Development of Improved HDAC6 Inhibitors as Pharmacological Therapy for Axonal Charcot-Marie-Tooth Disease. Neurotherapeutics, 14(2), 417–428. doi:10.1007/s13311-016-0501-z

Bradke, F., Fawcett, J. W., & Spira, M. E. (2012). Assembly of a new growth cone after axotomy: the precursor to axon regeneration. Nat Rev Neurosci, 13(3), 183–193. doi:10.1038/nrn3176

da Silva, T. F., Eira, J., Lopes, A. T., Malheiro, A. R., Sousa, V., Luoma, A.,… Brites, P. (2014). Peripheral nervous system plasmalogens regulate Schwann cell differentiation and myelination. J Clin Invest, 124(6), 2560–2570. doi:10.1172/JCI72063

Dan, W., Gao, N., Li, L., Zhu, J. X., Diao, L., Huang, J.,… Bao, L. (2018). alpha-Tubulin Acetylation Restricts Axon Overbranching by Dampening Microtubule Plus-End Dynamics in Neurons. Cereb Cortex, 28(9), 3332–3346. doi:10.1093/cercor/bhx225

Episkopou, V., Maeda, S., Nishiguchi, S., Shimada, K., Gaitanaris, G. A., Gottesman, M. E., & Robertson, E. J. (1993). Disruption of the transthyretin gene results in mice with depressed levels of plasma retinol and thyroid hormone. Proc Natl Acad Sci U S A, 90(6), 2375–2379. doi:10.1073/pnas.90.6.2375

Erturk, A., Hellal, F., Enes, J., & Bradke, F. (2007). Disorganized microtubules underlie the formation of retraction bulbs and the failure of axonal regeneration. J Neurosci, 27(34), 9169–9180. doi:10.1523/JNEUROSCI.0612-07.2007

Fleming, C. E., Mar, F. M., Franquinho, F., Saraiva, M. J., & Sousa, M. M. (2009). Transthyretin internalization by sensory neurons is megalin mediated and necessary for its neuritogenic activity. J Neurosci, 29(10), 3220–3232. doi:10.1523/JNEUROSCI.6012-08.2009

Fleming, C. E., Saraiva, M. J., & Sousa, M. M. (2007). Transthyretin enhances nerve regeneration. J Neurochem, 103(2), 831–839. doi:10.1111/j.1471-4159.2007.04828.x

Goldsteins, G., Andersson, K., Olofsson, A., Dacklin, I., Edvinsson, A., Baranov, V.,… Lundgren, E. (1997). Characterization of two highly amyloidogenic mutants of transthyretin. Biochemistry, 36(18), 5346–5352. doi:10.1021/bi961649c

Goodman, D. S. (1984). Vitamin A and retinoids in health and disease. N Engl J Med, 310(16), 1023–1031. doi:10.1056/NEJM198404193101605

Jochems, J., Boulden, J., Lee, B. G., Blendy, J. A., Jarpe, M., Mazitschek, R.,… Berton, O. (2014). Antidepressant-like properties of novel HDAC6-selective inhibitors with improved brain bioavailability. Neuropsychopharmacology, 39(2), 389–400. doi:10.1038/npp.2013.207

Kleele, T., Marinkovic, P., Williams, P. R., Stern, S., Weigand, E. E., Engerer, P.,… Misgeld, T. (2014). An assay to image neuronal microtubule dynamics in mice. Nat Commun, 5, 4827. doi:10.1038/ncomms5827

Komarova, Y., De Groot, C. O., Grigoriev, I., Gouveia, S. M., Munteanu, E. L., Schober, J. M.,… Akhmanova, A. (2009). Mammalian end binding proteins control persistent microtubule growth. J Cell Biol, 184(5), 691–706. doi:10.1083/jcb.200807179

Liz, M. A., Mar, F. M., Santos, T. E., Pimentel, H. I., Marques, A. M., Morgado, M. M.,… Sousa, M. M. (2014). Neuronal deletion of GSK3beta increases microtubule speed in the growth cone and enhances axon regeneration via CRMP-2 and independently of MAP1B and CLASP2. BMC Biol, 12, 47. doi:10.1186/1741-7007-12-47

Morley, S. J., Qi, Y., Iovino, L., Andolfi, L., Guo, D., Kalebic, N.,… Heppenstall, P. A. (2016). Acetylated tubulin is essential for touch sensation in mice. Elife, 5:e20813. doi:10.7554/eLife.20813

Moutin, M. J., Bosc, C., Peris, L., & Andrieux, A. (2020). Tubulin post-translational modifications control neuronal development and functions. Dev Neurobiol, 00, 1–20. doi:10.1002/dneu.22774

Plante-Bordeneuve, V., & Said, G. (2011). Familial amyloid polyneuropathy. Lancet Neurol, 10(12), 1086–1097. doi:10.1016/S1474-4422(11)70246-0

Rivieccio, M. A., Brochier, C., Willis, D. E., Walker, B. A., D’Annibale, M. A., McLaughlin, K.,… Langley, B. (2009). HDAC6 is a target for protection and regeneration following injury in the nervous system. Proc Natl Acad Sci U S A, 106(46), 19599–19604. doi:10.1073/pnas.0907935106

Silva, C. S., Eira, J., Ribeiro, C. A., Oliveira, A., Sousa, M. M., Cardoso, I., & Liz, M. A. (2017). Transthyretin neuroprotection in Alzheimer’s disease is dependent on proteolysis. Neurobiol Aging, 59, 10–14. doi:10.1016/j.neurobiolaging.2017.07.002

Woeber, K. A., & Ingbar, S. H. (1968). The contribution of thyroxine-binding prealbumin to the binding of thyroxine in human serum, as assessed by immunoadsorption. J Clin Invest, 47(7), 1710–1721. doi:10.1172/JCI105861

Xu, Z., Schaedel, L., Portran, D., Aguilar, A., Gaillard, J., Marinkovich, M. P.,… Nachury, M. V. (2017). Microtubules acquire resistance from mechanical breakage through intralumenal acetylation. Science, 356(6335), 328–332. doi:10.1126/science.aai8764

Yan, C., Wang, F., Peng, Y., Williams, C. R., Jenkins, B., Wildonger, J.,… Parrish, J. Z. (2018). Microtubule Acetylation Is Required for Mechanosensation in Drosophila. Cell Rep, 25(4), 1051–1065 e1056. doi:10.1016/j.celrep.2018.09.075

Zhang, Q., Fishel, E., Bertroche, T., & Dixit, R. (2013). Microtubule severing at crossover sites by katanin generates ordered cortical microtubule arrays in Arabidopsis. Curr Biol, 23(21), 2191–2195. doi:10.1016/j.cub.2013.09.018

